# Identification of Genetically Important Individuals of the Rediscovered Floreana Galápagos Giant Tortoise (*Chelonoidis elephantopus*) Provide Founders for Species Restoration Program

**DOI:** 10.1101/143131

**Authors:** Joshua M. Miller, Maud C. Quinzin, Nikos Poulakakis, James P. Gibbs, Luciano B. Beheregaray, Ryan C. Garrick, Michael A. Russello, Claudio Ciofi, Danielle L. Edwards, Elizabeth A. Hunter, Washington Tapia, Danny Rueda, Jorge Carrión, Andrés A. Valdivieso, Adalgisa Caccone

**Affiliations:** Department of Ecology and Evolutionary Biology, Yale University, 21 Sachem St. New Haven, Connecticut, 06520, United States of America; Department of Biology, School of Sciences and Engineering, University of Crete, Vasilika Vouton, Gr-71300, Heraklio, Crete, Greece; Natural History Museum of Crete, School of Sciences and Engineering, University of Crete, Knossos Av., GR-71409, Heraklio, Crete, Greece; College of Environmental Science & Forestry, State University of New York, Syracuse, New York, 13210, United States of America; Molecular Ecology Lab, School of Biological Sciences, Flinders University, GPO Box 2100, Adelaide, SA, 5001, Australia; Department of Biology, University of Mississippi, Oxford, Mississippi, 38677, United States of America; Department of Biology, University of British Columbia, Okanagan Campus, Kelowna, BC V1V 1V7, Canada; Department Biology, University of Florence, 50019 Sesto Fiorentino (FI), Italy; Life and Environmental Sciences, University of California, Merced, 5200 N Lake Rd, Merced, California, 95343, United States of America; Department of Natural Resources and Environmental Science, University of Nevada – Reno, Max Fleischmann Agricultural Building, Reno NV 89557 USA; Galapagos Conservancy, Fairfax, Virginia, 22030, United States of America; Galápagos National Park Directorate, Puerto Ayora, Galápagos, Ecuador

## Abstract

Species are being lost at an unprecedented rate due to human-driven environmental changes. The cases in which species declared extinct can be revived are rare. However, here we report that a remote volcano in the Galápagos Islands hosts many giant tortoises with high ancestry from a species previously declared as extinct: *Chelonoidis elephantopus* or the Floreana tortoise. Of 150 individuals with distinctive morphology sampled from the volcano, genetic analyses revealed that 65 had *C. elephantopus* ancestry and thirty-two were translocated from the volcano’s slopes to a captive breeding center. A genetically informed captive breeding program now being initiated will, over the next decades, return *C. elephantopu*s tortoises to Floreana Island to serve as engineers of the island’s ecosystems. Ironically, it was the haphazard translocations by mariners killing tortoises for food centuries ago that created the unique opportunity to revive this “lost” species today.

## Introduction

Human activities have generated an extreme and rapid loss of biodiversity^1^. Many actions have been undertaken to prevent species extinctions, including creating laws to protect endangered species and critical habitats^2^, translocations of individuals among populations or into new habitats^3,4^, and *ex situ* management including captive breeding^5^. For many species, such actions come too late to facilitate recovery^6–8^. Although ‘de-extinction’ using laboratory techniques is currently being debated^9^, such methods are viable only for some taxa and will generate significant anticipated, and unanticipated, risks^10,11^. Generally, extinction is final and cases where lost species can be revived will be extremely rare.

Despite their insularity, even remote oceanic islands are not exempt from rapid anthropogenic changes. For example, the ecosystems of the Galápagos Islands, located ∼900 kilometers off the Pacific coast of Ecuador, have been degraded by human activities since the archipelago was discovered in 1535. The resulting loss of biodiversity, including many endemic species, has been chiefly due to the introduction of non-native species^12,13^. In response, concerted efforts have been taken to restore ecosystems on the islands, including removal of introduced pests^14,15^ and population restoration via captive breeding and repatriation of threatened native species (e.g. ^16,17^).

Galápagos giant tortoises (*Chelonoidis* spp.) are flagship species for ongoing restoration efforts^18^ in this archipelago and play an important functional role as mega-herbivores in the islands’ ecosystems^19^. Galápagos giant tortoises can be classified into 15 species based on genetic data^20^. Generally, there is a single species per island with two exceptions: Isabela Island that has a different endemic species associated with each of its five volcanoes, and Santa Cruz Island that contains two species, one recently described^20^ (Figure 1a). The 15 species exhibit two general carapace shapes: "domed," a rounded cupola-like form (Figure 1c), and a "saddle-backed" form, with a high anterior opening creating the shape of a saddle (Figure 1d). Five of the 15 species have the saddle-backed morphology: *C. elephantopus* (Floreana), *C. hoodensis* (Española), *C. abingdoni* (Pinta), *C. ephippium* (Pinzón), and *C. chathamensis* (San Cristóbal; Figure 1a).

**Figure 1:**
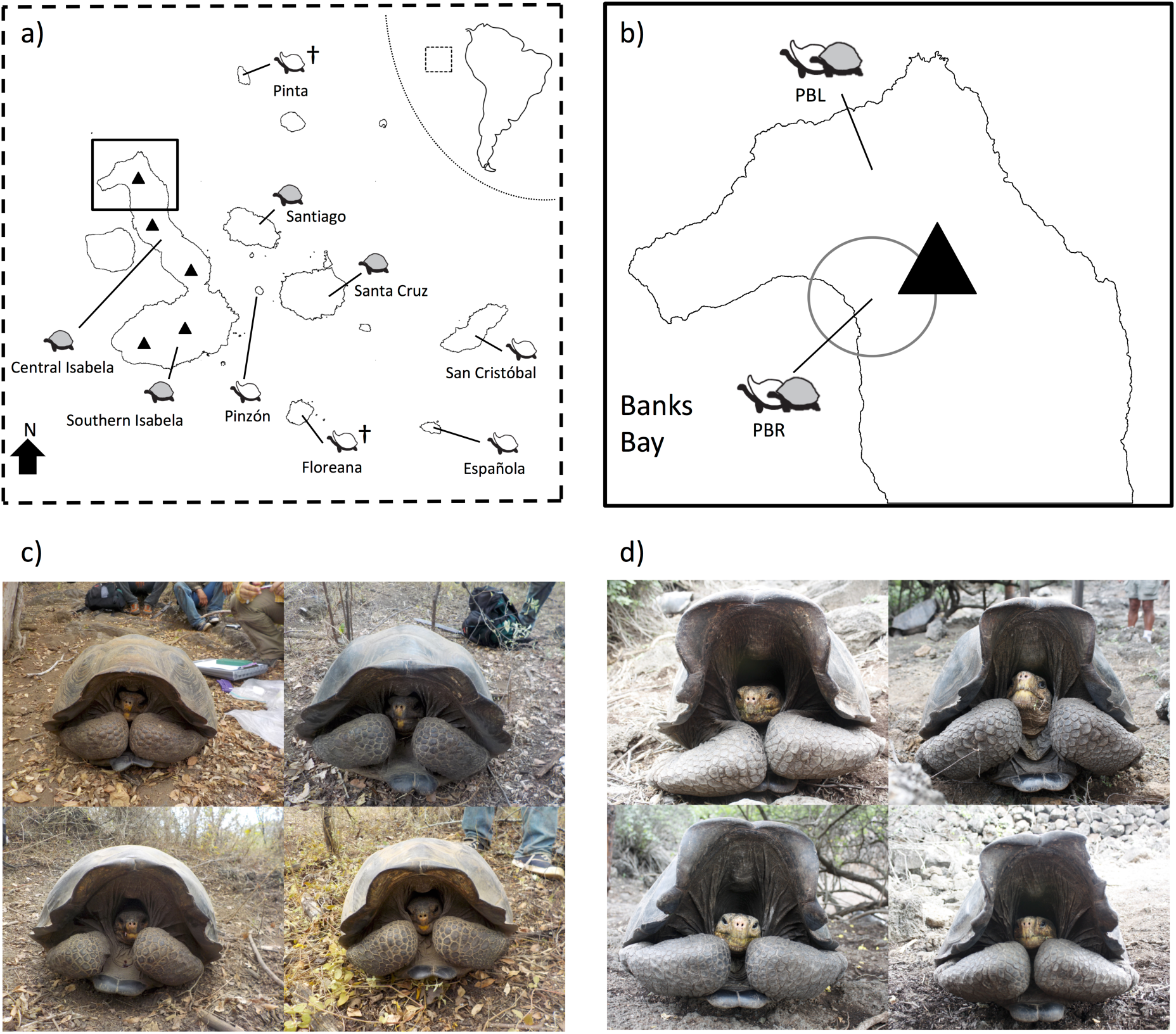
Distribution of tortoises among Galápagos Islands and representative photos of tortoise carapace morphology. A) Map of the distribution of tortoises among Galápagos Islands along with cartoons indicating carapace morphology for each. Light grey shading indicates domed morphology, unshaded indicates saddle-backed. Extinct species are noted with †. B) Larger view of Volcano Wolf on northern Isabela Island. The circle indicates the approximate field location of the current study. Examples of Galápagos giant tortoises with domed (C) saddle-backed (D) morphology. Maps were created with R (version 3.2.4^51^) using the raster package (version 2.5-8^52^).

Over the past three centuries, all giant tortoise populations experienced a ∼90% decline^21^, having been killed mostly for food and oil by whalers, sealers, buccaneers, and early colonists^22,23^. Four species have been declared Extinct^24^, including two of the five saddle-backed species: *C. elephantopus* from Floreana Island and, most recently, *C. abingdoni,* from Pinta Island. The latter species was represented by a single individual, Lonesome George, until his death in 2012.

Surprisingly, recent research found living, wild tortoises with genetic ancestry from two of the extinct saddle-backed species, *C. elephantopus* and *C. abingdoni* (hereafter referred to as the Floreana and Pinta tortoises, respectively) outside their native range^25–28^. These individuals, likely the descendants of tortoises translocated among islands by mariners^22,23,29^, were discovered on the remote Volcano Wolf on Isabela Island (Figure 1a,b). Among >1600 tortoises sampled from Volcano Wolf in 2008 during an exploratory expedition, 105 individuals were admixed between the locally endemic species, *C. becki*, and the Floreana (*n* = 86)^5^ or Pinta (*n* = 17)^4^ species. The majority of these genetically admixed individuals were found on Volcano Wolf’s western slopes, facing Banks Bay (also known as Puerto Bravo, PBR: Figure 1b), with a smaller number located on the volcano’s northwestern slopes near Piedras Blancas (PBL)^25,26^. No purebred individuals of either of the two non-native species were found in 2008, but genetic simulations and the young age of some mixed ancestry individuals indicated that purebred Floreana and Pinta tortoises might still be present on Volcano Wolf^25,26^.

Here we build on this previous work. In November 2015 we mounted a 10-day-long search involving ∼70 field personnel combined with helicopter- and ship-support. Focusing our search, we explored zones of Volcano Wolf most likely to contain individuals with ancestry from these two “lost” species (Figure 1b) and restricted genetic sampling to those individuals with saddle-backed morphology among the thousands of locally endemic domed *C. becki*. We then assigned ancestry to these tortoises using reference databases containing both extant and extinct species that have previously been used to assign ancestry to tortoises in the wild and captivity^25–31^. Based on these assignments, we determined suitability of the tortoises for a genetically informed captive breeding program aimed at reintroducing these key ecosystem engineers to their native island.

## Results & Discussion

In total, we encountered 144 individuals with saddle-backed morphology. Of those, 112 were released after taking blood samples, and 32 with pronounced saddle-backed morphology were transported to the Galápagos National Park Service’s captive tortoise breeding facility on Santa Cruz Island^32–34^. We assigned ancestry to all 144 of these individuals along with six saddle-backed tortoises known to have Floreana ancestry^35^ already residing at the breeding center using information from ∼700-bp of mitochondrial DNA sequence and diploid genotypes from 12 nuclear microsatellite loci. These loci have previously been shown to accurately assign individuals to tortoise species^25–31^.

Thirty-five of the 150 individuals analyzed had a mitochondrial DNA haplotype diagnostic of the Floreana species (Supplementary Figure S1). The remaining individuals either had haplotypes diagnostic of the Critically Endangered Española Island species (*n* = 70), haplotypes shared between the Santiago Island and Volcano Wolf species (*n* = 44), or a haplotype shared between the species from San Cristóbal and Santa Cruz Islands (*n* = 1). The proportions of haplotypes associated with the Española and Floreana Island species reported here (46% and 23%, respectively) are substantially higher than previously detected among individuals from Volcano Wolf^25,26^ (5% and 2%). This is likely due to our targeting of saddle-backed individuals in 2015 versus sampling broadly during previous surveys^25–28^.

Bayesian clustering analyses of microsatellite genotypic data using the method implemented in STRUCTURE^36^ revealed that 127 of the 150 tortoises sampled have ancestry assignments (Q-values) to the extinct Floreana species (average Q-value ± SD = 0.87 ± 0.21; range 0.16–0.99; Figure 2; Supplementary Figures S4-S6). Twenty-three individuals did not show evidence of Floreana ancestry, being assigned to the two genetically distinct populations (PBL and PBR) of *C. becki*, the endemic Volcano Wolf species. Of those individuals with Floreana Q-values, 30 had Floreana mitochondrial haplotypes (Figure 2).

**Figure 2:**
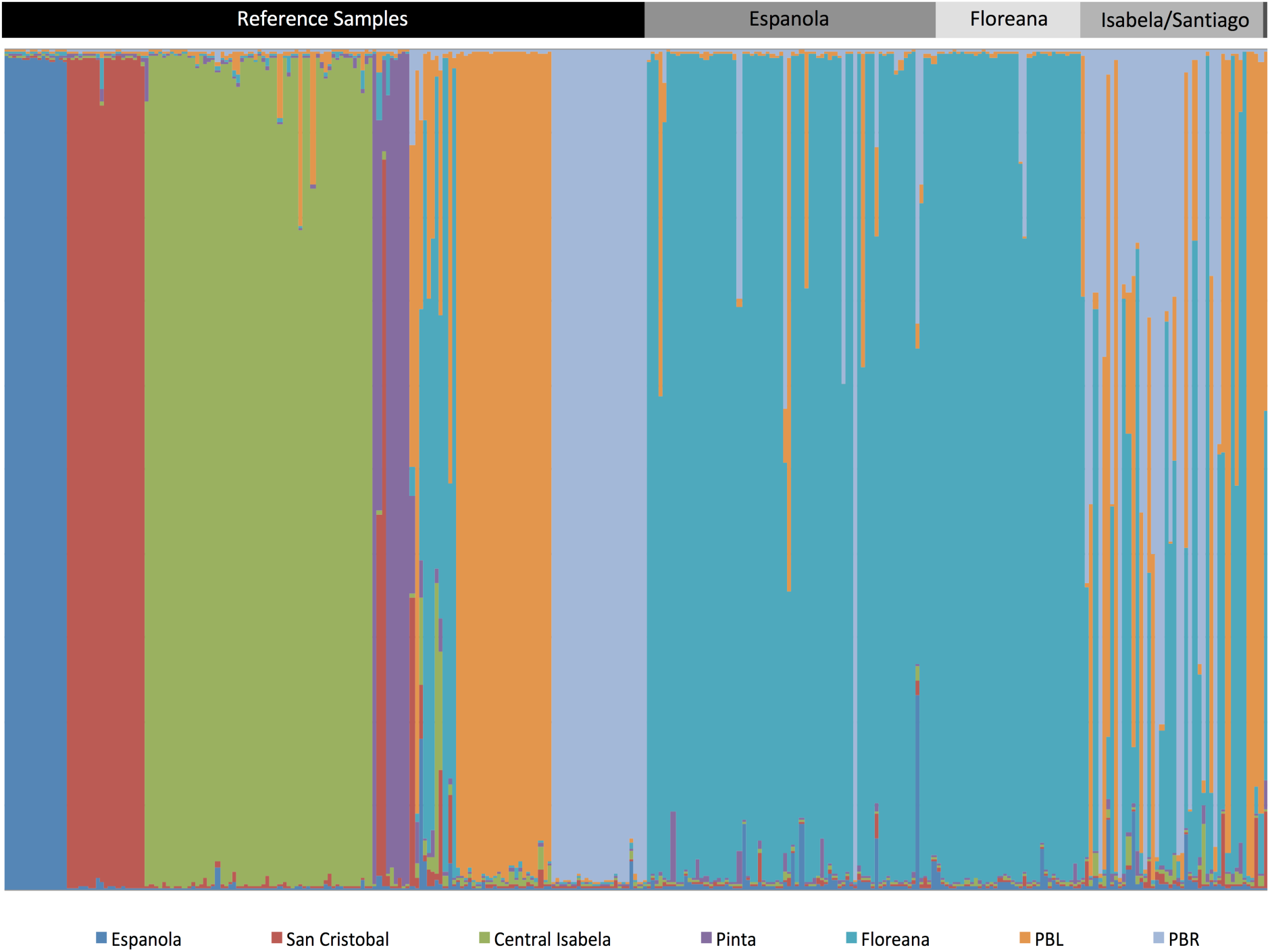
STRUCTURE^36,46^ plot for 155 reference samples along with the 150 newly collected individuals. Each individual is represented as a vertical bar, with colors denoting the different genetic clusters, as indicted below. The proportion of color in a bar is equal to the ancestry (Q-value) to a given cluster. Reference samples are those under the black horizontal bar, the newly collected samples are those under the grey horizontal bar with shades and labels corresponding to mitochondrial lineage (the final sample on the far right has a haplotype shared by the Islands of San Cristóbal and Santa Cruz).

We conducted additional assignment tests using genotypes generated by simulated matings within and among four possible parental lineages (PBL, PBR, Española Island, and Floreana Island). Use of simulated individuals in assignment tests has been suggested to improve accuracy and efficiency when distinguishing hybrid individuals^37^. Q-values from STRUCTURE for the simulated hybrids were in the range expected for each “class” of hybrid (Supplementary Figure S4), indicating we have the ability to identify individuals with Floreana ancestry across various levels of admixture.

Analyses with the program GeneClass2 version 2.0^38^ identified two tortoises with strong assignment to the Floreana species, being classified as either purebreds or backcrosses between a F_1_ × purebred Floreana tortoise (Supplementary Table S2). This program also identified additional 63 tortoises that were assigned to categories with Floreana ancestry. Further analysis of these individuals found a large number of F_1_ hybrids and backcrosses both when re-running STRUCTURE (Q-values between 0.40–0.77; Supplementary Table S2), as well as when we used NEWHYBRIDS version 1.1^39^ (Table 1) and discriminant analysis of principal components (DAPC; Figure 3, Table 1). Admixture between the saddle-backed species from Floreana and Española Islands (*n* = 43) was more common than between Floreana and either of the two endemic domed *C. becki* populations (PBL and PBR; *n* = 22), indicative of positive assortative mating between the two saddle-backed species.

**Figure 3:**
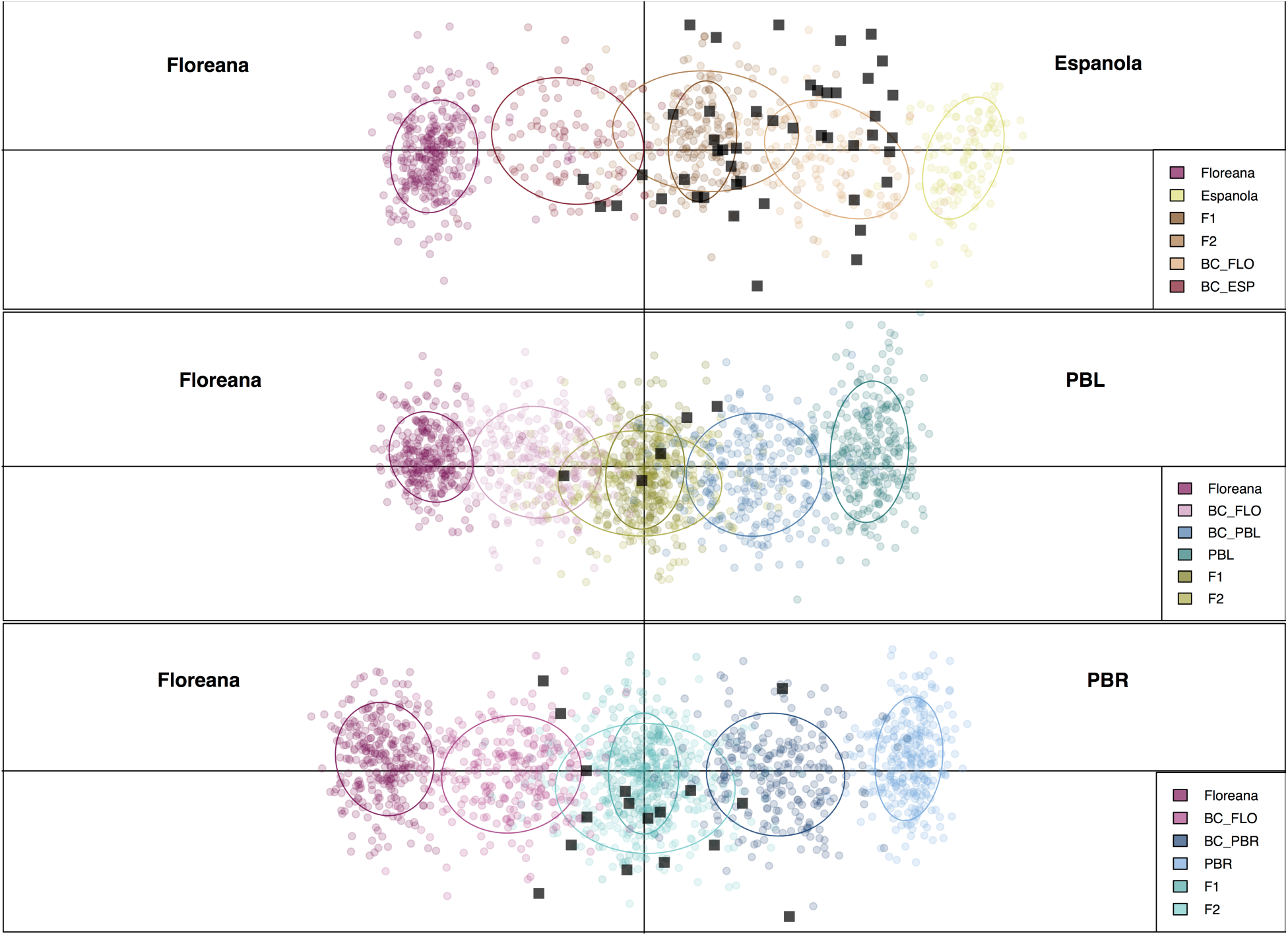
Scatterplot of the first two principal components of DAPC showing assignment of unknown individuals to ancestry categories. The rows show pairwise combinations of parental populations. Top: Española × Floreana; middle: Floreana × PBL; bottom: Floreana × PBR. Parental populations are positioned at the ends of each plot with color coded first generation (F_1_), second generation (F_2_), and each backcross (BC) individuals between them. Ellipses encompass ∼67% of the cloud of points for each group. These genotypes were used as a training set to define the discriminant functions and optimal number of PCs to retain. The Volcano Wolf tortoises with mixed ancestry are identified by black squares where placement is based on the previously defined discriminant functions.

**Table 1:** Number of individuals assigned to ancestry categories by DAPC (A) and NEWHYBRIDS (B)

|  |  | BC to<br>ESP | BC to<br>FLO | BC to<br>PBL | BC to<br>PBR | FLO/ESP<br>F<sub>1</sub> or F<sub>2</sub> | FLO/PBL<br>F<sub>1</sub> or F<sub>2</sub> | FLO/PBR<br>F<sub>1</sub> or F<sub>2</sub> | Total |
| --- | --- | --- | --- | --- | --- | --- | --- | --- | --- |
| Location |  |  |  |  |  |  |  |  |  |
|  | PNG | 9 | 2 | 0 | 0 | 8 | 1 | 3 | 23 |
|  | VW | 13 | 5 | 1 | 3 | 12 | 2 | 8 | 44 |
| Total |  | 22 | 7 | 1 | 3 | 20 | 3 | 11 | 67 |
| Location |  |  |  |  |  |  |  |  |  |
|  | PNG | 0 | 12 | 0 | 0 | 4 | 1 | 2 | 19 |
|  | VW | 0 | 7 | 0 | 0 | 17 | 4 | 8 | 36 |
| Total |  | 0 | 19 | 0 | 0 | 21 | 5 | 10 | 55 |

The current location of individuals is listed, as Galápagos National Park Breeding Center (PNG) or Volcano Wolf (VW). Abbreviations are: BC = backcross, F<sub>1</sub> = first generation mating, F<sub>2</sub> = second generation mating (F<sub>1</sub> x F<sub>1</sub>), ESP = Española, and FLO = Floreana. Note that the total number of samples assigned differs between the two methods as some individuals did not exceed the posterior probability threshold for classification in NEWHYBRIDS (see Supplementary Methods).

A critical factor when planning a captive breeding program is having accurate ancestry assignments. To examine the precision and accuracy of our ancestry assignments we conducted additional analyses in STRUCTURE. First, jackknifing the loci to test robustness of Q-values to reductions in the number of markers, and using the ANCESTDIST option in STRUCTURE, which collects information about the distribution of Q-values for each individual in our dataset. Jackknifing showed that individual Q-values were robust to reductions in the number of markers: average variation in individual Q-values was 0.027 (SD=0.033, range <0.001-0.149; Supplementary Figure S5). Second, use of the ANCESTDIST option highlighted that, as expected, for hybrid individuals there is uncertainty around the specific estimate of Q-values (range 0.002-0.484; Supplementary Figure S6). Inclusion of the simulated individual in our STRUCTURE run led to a general decrease in the magnitude of Q-values observed (Supplementary Table S2 and Supplementary Figure S7). Taken together, these analyses suggest that Q-values should not be taken as direct measure of proportional ancestry, but that our markers are powerful in detecting the presence of ancestry from the extinct Floreana species.

Neither mitochondrial DNA nor microsatellite data identified individuals with ancestry from the Pinta Island species. However, a previous genetic estimate suggested that only 60–70 tortoises with Pinta ancestry are present on Volcano Wolf^25^, whereas capture-mark-recapture methods employed during our expedition estimated that a total of ∼5,000–6,000 tortoises occurred in the area searched. Moreover, despite our substantial search effort, we explored only ∼26% of the total tortoise-occupied range on the volcano (J. Gibbs unpublished data). Therefore, it is possible that individuals with Pinta ancestry still live on Volcano Wolf, but went undetected in 2015.

Of the 38 individuals currently housed in the captive breeding center, 23 tortoises (9 males and 14 females) were found to have nuclear ancestry from Floreana across multiple assignment methods (Table 1) with Q-values from STRUCTURE ranging between 0.44–0.77, when simulated individuals were also included in the analyses. In addition, 12 of these 23 individuals have mitochondrial haplotypes from the Floreana species (Supplementary Table S2). Together, these 23 individuals now form the core of a genetically-informed captive breeding program aimed at repatriation of tortoises to Floreana Island. For the purposes of the breeding program, all 23 genetically important individuals will be included, regardless of mitochondrial lineage, in order to capture and maintain as much nuclear genetic diversity from the Floreana species as possible. The program is modeled after another one developed for the Española species initiated with only 15 founders. Over 50 years, the Española tortoise program generated >2000 repatriates with nearly 1,000 surviving tortoises now reproducing independently on their native island^17,40,41^.

A key attribute for success of such breeding programs is that the founding individuals are unrelated, as high relatedness can lead to inbreeding depression. Accordingly, we examined relatedness among the 23 individuals with Floreana ancestry in the breeding center. The analysis showed that most individuals were unrelated (average Queller and Goodnight’s^42^ relatedness = -0.04, range -0.58–0.63; Supplementary Figure 3). Although the 23 tortoises in captivity represent a promising founding population, 44 tortoises identified during the 2015 expedition, but left on Volcano Wolf, are also good candidates for the Floreana breeding and repatriation program (Table 1). These individuals, once re-located on Volcano Wolf, could be incorporated into the breeding program to further expand the genetic diversity of the founder population.

The Floreana tortoise breeding program will be designed to maximize founder contributions and *C. elephantopus* genome representation in the resulting progeny, while promoting *in situ* population growth and minimizing costs to the Galápagos National Park Directorate. Depending on the goals and priorities of Park decision-makers, complete genome recovery may not be reached before releases of offspring begin. However, the high proportion of Floreana ancestry and low relatedness evident in the current breeding individuals indicate that the 23 founding individuals and their resulting progeny will provide a good starting point for restoring the species.

Our discovery raises the possibility that the extinct Floreana species could be revived. In this case, tortoises with Floreana ancestry are living ‘genomic archives’ that retain the evolutionary legacy of the extinct species, removing the need for the cloning methods that have been proposed to bring back extinct species^43^. The Floreana tortoise breeding program is anticipated to generate thousands of offspring over the next few decades. When repatriated to Floreana Island, these tortoises can once again play their critical role as ecosystem engineers^19^. In addition, giant tortoises are a major tourist attraction in Galápagos^44^; tortoise restoration on Floreana Island should create new economic opportunities for the island’s few human residents. Ironically, the opportunity to revive this “lost species” today was created by the same early visitors to the archipelago whose activities imperiled most giant tortoise species and drove some into outright extinction.

## Materials and Methods

### Sampling and lab methods

All samples were collected under CITES permit 15US209142/9, Galápagos Park Permit PC-75-16, and in accordance with Yale Institutional Animal Care and Use Committee (IACUC) permit number 2016-10825. Samples were collected over a 10-day expedition in November 2015 by a team of 36 researchers plus Galápagos National Park rangers. Groups of 3–4 searchers were assigned distinct areas on Volcano Wolf, collectively totaling ∼36 km^2^. The search area was chosen based on a previous survey of tortoises on Volcano Wolf, which indicated that individuals with Pinta and Floreana ancestry were found to be in their highest densities on the western slopes of Volcano Wolf^25^. For all tortoises encountered, sex, age, and GPS coordinates were recorded; for saddle-backed individuals, photographs were taken, blood samples were collected for DNA analysis, and each individual was injected with a passive integrated transponder (PIT) tag under the skin for identification in the future. In cases where blood was taken, ∼2 ml of blood was collected from the brachial vein of one of the front legs of the tortoise and preserved in a lysis buffer containing 0.1 M Tris buffer, 0.1 M EDTA, 0.2 m NaCl, and 1% SDS, pH 8.0. All tortoises were uniquely marked with paint when first encountered. Starting on the sixth day, we re-searched areas and recorded whether any individual had been previously encountered for the purposes of a capture-mark-recapture estimate of population size. In total, 1,333 tortoises were encountered, of which 144 had saddle-backed morphology. The 32 tortoises removed from Volcano Wolf were initially carried in nets by helicopter from the flanks of the volcano to the expedition ship anchored in Puerto Bravo harbor, and then transported to the Galápagos National Park captive tortoise breeding facility on Santa Cruz Island.

DNA was extracted from 150 blood samples using Qiagen blood and tissue extraction kits. These samples included the 144 saddle-backed individuals mentioned above along with six individuals already housed at the Galápagos National Park Breeding Center on Santa Cruz Island that were previously identified to have Floreana ancestry^35^. All samples were sequenced at ∼700bp of the mitochondrial DNA control and were genotyped at 12 dinucleotide microsatellites, using previously developed protocols (detailed procedures in Supplementary Methods).

### Ancestry assignment

For ancestry assignment based on mitochondrial DNA, the new sequences were aligned to a reference database of 123 previously observed haplotypes, representing all extant and extinct species^29^. Ancestry was assigned by determining shared haplotypes using the program TCS version 1.21^45^. Evolutionary relationships were viewed with the program Network version 5.0 (fluxus-engineering.com; Supplementary Figure 1).

For ancestry assignment based on microsatellite genotypic data, we took a two-step process using STRUCTURE version 2.3.4^36,46^. First, we ran STRUCTURE with a reference dataset of 277 samples including all extant and extinct species to confirm the number of expected genetic clusters (*K*) present in the archipelago (for full parameters see Supplementary Methods). In this case, the optimal *K* was 12, which corresponds to previously described results^25,26^ (Supplementary Figure 2). Second, with archipelago-wide *K* established, we reduced the reference dataset to the seven clusters previously found to be on Volcano Wolf^29^: Española, San Cristóbal, Central Isabela (La Cazuela, Volcano Alcedo, and Volcano Darwin), Floreana, Pinta, and the two genetically distinct endemic Volcano Wolf populations, PBL, and PBR. To this reduced dataset (155 reference samples in total), the new samples were added and we set *K* to 7, leaving all other parameters unchanged.

To further quantify the genetic ancestry of individuals, assignment tests were undertaken using three additional methods. Prior to implementing these methods, we expanded the reference database to include simulated genotypes. These genotypes were simulated using HYBRIDLAB version 1.0^47^, and corresponded to individuals arising from crosses within and among the genetically distinct *C. becki* populations on Volcano Wolf (PBL and PBR), and the two saddle-backed species from Española and Floreana (see Supplementary Methods). These four lineages were chosen due to their high prevalence in the mitochondrial DNA-based assignments and their contribution to microsatellite genotypes in admixed individuals. Simulated individuals represented explicit ancestry categories: pure parental populations, first-generation (F_1_) crosses, second-generation (F_2_) crosses, and backcrosses (i.e., matings between F_1_’s and their respective purebred lineages).

With these new genotypes added into the reference set, we first used GeneClass version 2.038 to calculate the probability that an individual’s genotype assigns to a population. Second, we used NEWHYBRIDS version 1.1^39^ to compute the posterior probability of various hybrid classes for each individual. In this case, only four pairwise combinations of parental populations were considered to focus on identifying potential hybrids involving Floreana ancestry: 1) Española × Floreana, 2) PBL × Floreana, 3) Floreana × PBR, and 4) Española × PBR. Third, we assigned ancestry using a multivariate approach, DAPC, as implemented in the R package *adegenet* version 2.0.1^48,49^. In this case, the simulated genotypes from crossess among Floreana, Española, PBL and PBR were initially used as a training dataset to define the principal components and discriminant functions. The empirical genotypes for putative hybrids were then transformed with principal components analysis (PCA) based on the centering and scaling of the training set, and positioned onto the discriminant functions. The NEWHYBRIDS and DAPC analyses were carried out only on individuals with ancestry from Floreana based on the results from the GeneClass analyses (Supplementary Table S2). Finally, we re-ran STRUCTURE using the simulated crosses as the reference populations.

Precision and accuracy of ancestry estimates were tested with two additional sets of STRUCTURE analyses. First, we jackknifed our loci, sequentially removing one locus from the dataset and rerunning STRUCTURE. Second, we used the ANCESTDIST option within STRUCTURE. See Supplementary Materials for full methods.

### Relatedness Analysis

For the 23 individuals in the breeding center with Floreana ancestry, we calculated pairwise relatedness using the estimator of Queller and Goodnight^42^. This was the highest ranked of eight tested relatedness estimators tested by the program irelr^50^ which considers a composite score that incorporates estimates of bias, variance, skewness, and kurtosis. The empirical distribution of relatedness values was compared to the distributions of 10,000 simulated pairs of individuals for each of four relatedness categories (unrelated, half sibs, full sibs, and parent– offspring). The pairwise relatedness distribution of both the empirical and simulated data were calculated using irelr^50^.

## Acknowledgements

This work was enabled by substantial logistical and personnel support of the Galápagos National Park Directorate through its dedication to science-based conservation of giant tortoises via the “Giant Tortoise Restoration Initiative” undertaken in conjunction with the Galapagos Conservancy. Expedition expenses and laboratory analyses were also supported by grants from the Galapagos Conservancy, the Mohamed bin Zayed Species Conservation Fund, National Geographic Society, and the Oak Foundation to AC, the US National Science Foundation (DEB 1258062) to JG, the Belgian American Educational Foundation to MQ, and the Yale Institute for Biospheric Studies. C. Gottsegen helped with genetic data collection, A. Antoniou provided assistance with DAPC analyses. L. Boitani, L. Cayot, C. Parker, and A. Rhodin helped procure funds and provided encouragement. J. Flanagan provided critical veterinary support. We would also like to thank C. Cullingham for insightful comments on the manuscript.

## Author Contributions

JMM, NP, LB, CC, EAH, WT, DR, JC, JPG and AC assisted with field logistics and sample collection. JMM, MCQ, and AV conducted laboratory analyses. JMM, MCQ, and NP conducted data analyses. LBB, RCG, MAR, and DLE provided technical guidance and conceptual advice for data analyses. JMM drafted the manuscript with assistance from MCQ, LBB, MAR, JPG, and AC. JMM, MCQ, NP, LBB, RCG, MAR, CC, DLE, EAH, WT, JPG, and AC discussed the results and implications and commented on the manuscript at all stages.

## Additional Information

Microsatellite genotypes and mitochondrial sequences are available on a server hosted by the College of Environmental Science & Forestry, State University of New York, Syracuse (http://www.esf.edu/efb/gibbs/Miller_et_al_data_archive.zip). The authors have no financial conflicts of interest. Correspondence and requests for materials should be addressed to either.

